# The dynamic tumor ecosystem: how cell turnover and trade-offs affect cancer evolution

**DOI:** 10.1101/270900

**Authors:** Jill A. Gallaher, Joel Brown, Alexander R. A. Anderson

**Author notes:** These authors contributed equally (Joel Brown, Alexander R. A. Anderson).

## Abstract

Tumors are not static masses of cells but rather dynamic ecosystems where cancer cells experience constant turnover and evolve fitness-enhancing phenotypes. Selection for different phenotypes may vary with 1) the tumor niche (edge or core), 2) cell turnover rates, 3) the nature of the tradeoff between traits (proliferation vs migration), and 4) whether deaths occur in response to demographic or environmental stochasticity. In an agent based, spatially-explicit model, we observe how two traits (proliferation rate and migration speed) evolve under different trade-off conditions with different turnover rates. Migration rate is favored over proliferation at the tumor’s edge and vice-versa for the interior. Increasing cell turnover rates only slightly slows the growth of the tumor, but accelerates the rate of evolution for both proliferation and migration. The absence of a tradeoff favors ever higher values for proliferation and migration. A convex tradeoff tends to favor proliferation over migration while often promoting the coexistence of a generalist and specialist phenotype. A concave tradeoff slows the rate of evolution, and favors migration at low death rates and proliferation at higher death rates. Mortality via demographic stochasticity favors proliferation at the expense of migration; and vice-versa for environmental stochasticity. All of these factors and their interactions contribute to the ecology of the tumor, tumor heterogeneity, trait evolution, and phenotypic variation. While diverse, these effects may be predictable and empirically accessible.

## I. INTRODUCTION

Tumors are thought to consist of 3 major populations of cells: actively dividing, quiescent and necrotic. Under idealized environments, such as the experimental system of spheroids (1), a fast growing tumor becomes dense and quickly outgrows the supply of oxygen and nutrients. This gives rise to a layered tumor anatomy that consists of concentric regions encompassing the 3 populations (e.g. Fig. 1A). In real tumors, the geometry of these regions appears far more irregular and disordered (e.g. Fig. 1B), reflecting a the more complex and dynamic environment. Regardless, it is a tempting simplification to view the tumor edge as the place where tumor cells primarily divide rather than die, the interior as generally quiescent with few births and deaths, and the necrotic zone where tumor cells mostly die.

**Figure 1.**
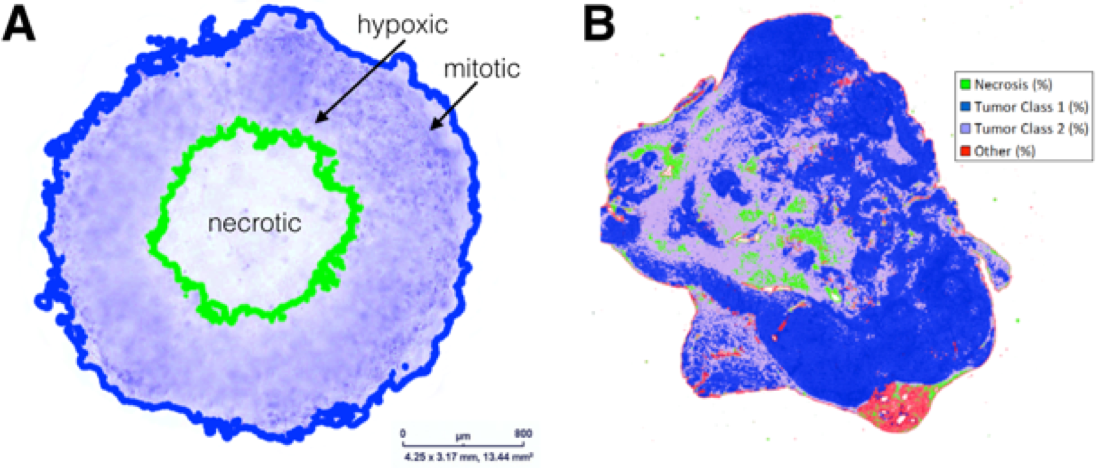
Comparing structures of tumor models and patient tumors. A) Tumor spheroid model. Edge detection algorithm finds inner necrotic (green) and outer proliferating (blue) edges. B) Digital pathology uses pattern recognition on histological sample from actual tumor. The proliferating, hypoxic and necrotic regions have the same broad structure but are more intermixed.

Such a perspective has led to models of tumor growth and evolution where tumor cells expand to occupy space, either explicitly (2–7) or implicitly (8,9), as different clonal lineages proliferate and expand at different rates. When these models include evolution, one can determine the properties of tumor cells that are favored by natural selection. Such is the case for models that examine the joint evolution of proliferation and migration (2,3,6). However, in the absence of cell turnover, such models can only show changes in the frequency of different clonal lineages while the replacement of less successful lineages by more successful ones is ignored.

In reality, the turnover of tumor cells via proliferation and cell death occurs constantly throughout the entirety of the tumor. Turnover rates may be high, perhaps as high as every 10 days for the interior of breast cancer tumors. A tumor that looks static with an unchanging volume might actually be very dynamic as proliferation and apoptosis occur in parallel throughout a tumor. High but balanced proliferation and death rates have been measured in some cancers (10–13). Furthermore, stimulatory factors from dying cells can cause compensatory proliferation of surviving cells (14), and an increased proliferation along with an increased death rate may suggest a more aggressive disease (12,13).

High turnover rates facilitate evolution by natural selection (15). This “struggle for existence” is seen in all organisms, and in cancer the cells have the capacity to produce more offspring than can possibly survive. Competition for space and resources limits cancer cell densities and population sizes. Limits to growth and cell turnover should select for genes and traits associated with proliferation rates and movement. All else equal, the cancer cell lineage with a higher proliferation rate will outcompete and replace one with a slower proliferation rate. However, higher proliferation rates will cause local crowding, limitations on resources, and other unfavorable conditions. Movement and migration away from such crowding should be favored. Even random migration can be favored by natural selection as a means of avoiding over-crowding (16). Such migration can be particularly favorable at the edge of the tumor, but even in the interior of a tumor, migration may move cells from more to less dense locales.

Many mutation models of cancer progression allow for unconstrained phenotypic improvement (2,3,5) or infer increased fitness through the number of passenger/driver mutations (17,18). Indeed, if both proliferation and migration enhance the fitness of cancer cells, then natural selection should favor higher rates for both. Such selection will continue to improve proliferation and migration rates simultaneously until a point is reached where there are tradeoffs (19–21). To improve proliferation rates further necessarily means sacrificing migration and vice-versa (22–24). In his seminal book on evolution of changing environments, Levins (1967) proposed that the shape of the tradeoff curve should influence the evolutionary outcome (25). A convex curve may favor a single population with a generalist phenotype whereas a concave curve may favor the coexistence of two specialist populations. Additionally, the specific shape of the trade-off curve can significantly affect the evolutionary trajectory towards this curve (26).

The pattern of cancer cell mortality across a tumor may represent just *demographic stochasticity* or it may include *environmental stochasticity* (27). The former happens when cell death is random and exhibits little temporal or spatial autocorrelations. Such patterns of mortality open up numerous but small opportunities for cell replacement. Environmental stochasticity happens when the sudden absence of nutrients or the accumulation of toxins causes wholesale death of the cells in some region of the tumor. This pattern of cell mortality creates fewer but much larger spaces for cell replacement. When regions are subject to catastrophic death (e.g., large or small temporary regions of necrosis) the distinctiveness of edge versus interior regions of a tumor are obscured, and the evolution of different combinations of proliferation and migration rates may be favored. Strictly demographic stochasticity should favor proliferation over migration and vice-versa for environmental stochasticity within the tumor.

In what follows, we develop a spatially explicit agent-based model of tumor growth that includes cell turnover at both the edge and the interior of the tumor. We use this model to explore the joint evolution of proliferation and migration rates by cancer cells in response to: 1) rates of cell turnover, 2) different shapes of the tradeoff curve, 3) and different mortality regimes.

## II. MATERIALS AND METHODS

The agent-based computational model was created to investigate evolving phenotypes under different constraints. The phenotype is defined as a combination of two traits: the proliferation rate and the migration speed of the cell. The simulation is initialized with one cell in the least aggressive state (lowest migration speed = 0 microns/h and slowest proliferation rate = 50h inter-mitotic time) centered in a 4mm circular tissue and updated every minute. Given the proliferation rate, when it is time for the cell to divide, it will split into two daughter cells, each taking traits within a range of the mother cell’s traits. The cells otherwise are allowed to move throughout a 2-dimensional space at a certain speed. Specifically, they follow a persistent random walk, designated from a normal distribution of persistence times (mean 80 min and standard deviation 40 minutes) and random turning angles. If the cells overlap in space or hit the boundary of the space (circle of diameter ~2.7mm), they change direction following a lossless collision. If the cells run out of space in their immediate neighborhood, they stop dividing and migrating.

**Figure 2.**
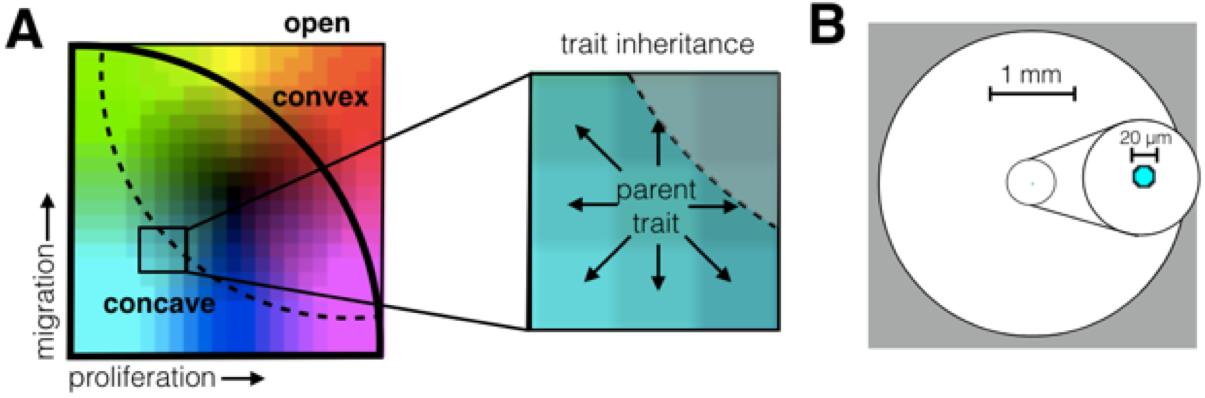
Model setup. A) Imposing trade-offs by bounding the phenotype space. When the whole space is open (thin solid line), all phenotypes are allowed. The convex (thick solid line) and concave (dashed line) bound the space as shown. The set of evolutionarily feasible traits lies within (fitness set) and on the tradeoff line (active edge). The trait value of a daughter cell can mutate up to one unit in any direction as long as it stays within the bounded region. B) A single cell with the smallest proliferation and migration rates centered in a 4-mm radial boundary initializes the simulation. Cell diameter is 20 μm.

### BOUNDING THE TRAIT SPACE

We limit the possible trait combinations according to i) an infinite improvement model, ii) a convex curve, and iii) a concave curve (see Fig. 2A). When a cell divides, new traits are determined giving each option (improve, stay the same, or diminish) the same weight. If the current trait is already on the boundary of trait space, then only options that respect this bound are considered and weighted equally. For the convex case, the forbidden region is created by making a circular arc from the two extreme values where fitness is greatest for each trait but worst for the other (i.e. point A with 10 h IMT and 0 microns/h and point B with 50 h IMT and 20 microns/h). The trait combinations with the fastest proliferation rates and fastest migration speeds are not allowed. For the concave case, the forbidden region cuts off this space as well, but goes even further. The circular arc is created with opposite concavity and through the points with one extreme value and the other a quarter along the range (i.e. point A with 10 h IMT and 5 microns/h and point B with 40 h IMT and 20 microns/h).

### CELL DEATH

Cell death either occurs randomly distributed or regionally clustered (catastrophic). The probability of death is split between these two types of death with either all random, all catastrophic, or 1:1 mix of random and catastrophic. Random death occurs with a given probability for every cell at every frame. When there are catastrophic death events, all cells within a confined circular region 500 μm in diameter, which is randomly placed, will die. The cells don’t automatically die but wait a randomly chosen period between 6-15 hours before being removed from the system. This is an estimate for how long it takes to go through apoptosis (28,29).

The probability of death for a single cell is once per week for the high death rate and once every two weeks for the low death rate. The actual death rate is variable because it depends on the number of cells at any time, but when the space is completely full (approximately 13,000 cells), around 2,000 cells are dying per day for the high death rate and 1,000 cells per day for the low death rate.

For the catastrophes, we need to ensure that the number of cells deaths on average is similar to the random death rate, because they happen at a population level at certain time points rather than to individuals. We define the probability of a catastrophic event *p*_cat_ based on the probability of death *p*_death_ and the time intervals *T*_cat_ between catastrophic events:

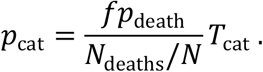

Here *f* is the fraction of deaths that are catastrophic, *N*_deaths_ is the number of cells that die from each catastrophic event, and *N* is the total number of cells. Setting the probability of a catastrophic event *p*_cat_ to 1, we can solve for *T*_cat_ to get the appropriate time between catastrophes:

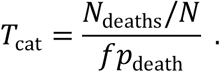

However, because the catastrophic death region will be spaced randomly there is a possibility that a new catastrophe will overlap with an old one before filling back in or will lie on an edge, so in general, there won’t be the same number of cells that die each time a catastrophe occurs. This can be accounted for if the time between events is changed each time based on the number of deaths from the previous event. If the number died previously is less than what would be given by *fp*_death_, then the numerator gets smaller, making a smaller time interval between events, and if the number that died is larger, then the next time interval will be larger. By adjusting after each event, we can compensate for this variation.

## III. RESULTS

Using an off-lattice agent-based model, we investigated how traits will evolve in response to space limitation and the continual turnover of cells. Figure 2A shows the trait space with respect to proliferation and migration, and Fig. 2B shows the 4mm diameter circular space available to the tumor.

Initially, we start with a single cell with the least aggressive phenotype: a long cell cycle time and a slow migration speed. Upon division, each daughter cell’s trait may change in one of three ways: it can inherit the same trait as the original cell, or via mutation its trait values for migration or proliferation rate can increase or decrease by a small value, so long as its trait values stay within the boundaries of what is evolutionarily feasible. Density-dependence and limits to population growth comes from local crowding. When a cell is completely surrounded by neighbors, we assume that it can neither move nor divide. More detail on how the carrying capacity is imposed in an off-lattice model can be found in Gallaher et al (3).

### IMPOSING A GO-OR-GROW TRADE-OFF SELECTS FOR MIGRATION DURING GROWTH

Evolutionarily, we placed limits on the set of feasible combinations of migration and proliferation. The boundary of this set represents the tradeoff between the two traits. In our simulations, we considered three forms of the tradeoff: open, convex, and concave boundary conditions (Fig. 2A). Under an open tradeoff, each trait can achieve a maximum value independent of the value of the other trait (no tradeoffs). Under a convex (or concave) tradeoff, the maximum feasible values for migration and proliferation occur along a curve that bows outwards (or inwards). Regardless of the shape of the boundary, natural selection should favor cancer cells with ever greater migration and proliferation rates until reaching the boundary edge. However, the shape of the tradeoff may influence both the evolutionary trajectory of the cancer cells, their evolutionary endpoint, and the diversity or variance of trait values among the cancer cells.

Ecologically, we first considered the case where there is no cell mortality. In this case, the population of cells will divide and migrate until the space is filled completely (see Fig. 3A for the spatial layout). In the absence of death, we see rings of cells with different phenotypes within the tumor. While natural selection favors cells with greater trait values, these trait values can only arise through successive cell divisions. The least aggressive cells, those with the lowest trait values, form the core (cyan color). Towards the outer edges, cells with more aggressive traits predominate at the periphery. Cells that mutate with higher proliferation rates can increase in frequency where space permits, and cells that mutate with higher migration rates can move into empty spaces where longer runs of proliferation are possible. Even as the whole population evolves, each step in this evolution leaves tree ring like layers in the tumor. With no cell death, the entire historical record in space and time is preserved.

Once the space has filled, the distribution of cancer cell phenotypes can be seen in Fig. 3B in the form of a density map. Color intensities as seen in Fig. 2 correspond to the relative frequency of phenotypes where white indicates an absence of cells with that phenotype. As expected, when the tradeoff boundary between migration and proliferation is open, the most frequent phenotypes exhibit both fast proliferation and fast migration. As the tradeoff boundary changes from open to convex to concave, we see natural selection favoring migration over proliferation. Contrary to expectation, the convex tradeoff boundary did not produce a generalist phenotype. Instead there is an apparent coexistence of two cell types: one with high migration but moderate proliferation, the other just the opposite. Also, contrary to expectations, a concave tradeoff boundary did not promote the coexistence of extreme phenotypes but instead, natural selection favored higher migration with little to no improvement in proliferation rates.

**Figure 3.**
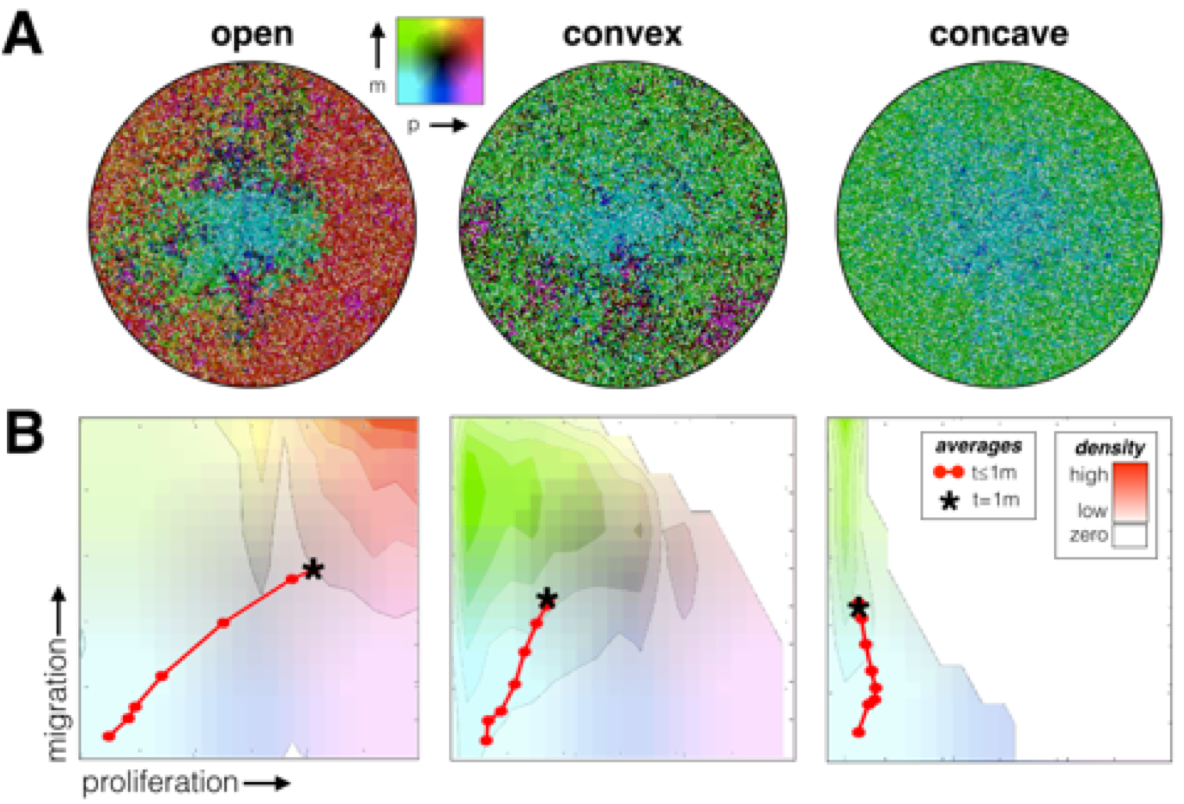
Joint evolution of migration and proliferation as influenced by three different tradeoff boundaries: open, convex, and concave. A) The spatial layout is shown and B) and the frequency of trait combinations is shown, where the red points and line mark the average trait values every 5 days. The background colors correspond to the density of cell traits after reaching capacity; Brightly colored areas correspond to high densities, and the completely white area contains no cells.

The sequence of red dots in Fig. 3B show the evolutionary trajectory over time, of the average values, of proliferation and migration rates within the simulated tumor. Each point gives the average phenotype in increments of 5 days until the space is filled. From the spacing of the dots, we see that an open tradeoff boundary produces rapid evolution, rapid space filling, and the highest level of average proliferation rates. A concave tradeoff boundary results in the slowest evolution, slowest space filling, and the lowest average proliferation rate. In going from open to convex to concave tradeoff boundaries, the phenotypes become less proliferative. Thus, they divide, evolve, and fill space more slowly.

### AN INCREASED DEATH RATE SELECTS FOR INCREASED PROLIFERATION

We examined the eco-evolutionary consequences of cell turnover by incorporating random cell death. Figure 4 shows the results when there is no death (top), and a low (middle) and high (bottom) random death rate. The spatial layout is shown in Fig. 4A, and a density map representing the frequency of trait combinations is shown in Fig. 4B for the 3-month time point. The evolutionary trajectory of the average trait values the population took for the first 3 months are overlaid on the density map, shown in red, while the black asterisk shows the average phenotype at 12 months.

With non-zero death rates, the phenotypic evolution has two apparent phases: the first occurs while space is relatively sparsely occupied, and the second occurs through cell turnover after the space has filled. During the first phase, phenotypic evolution follows a similar trajectory as the case when there is no death. However, as space fills, selection favors faster proliferation rates.

**Figure 4.**
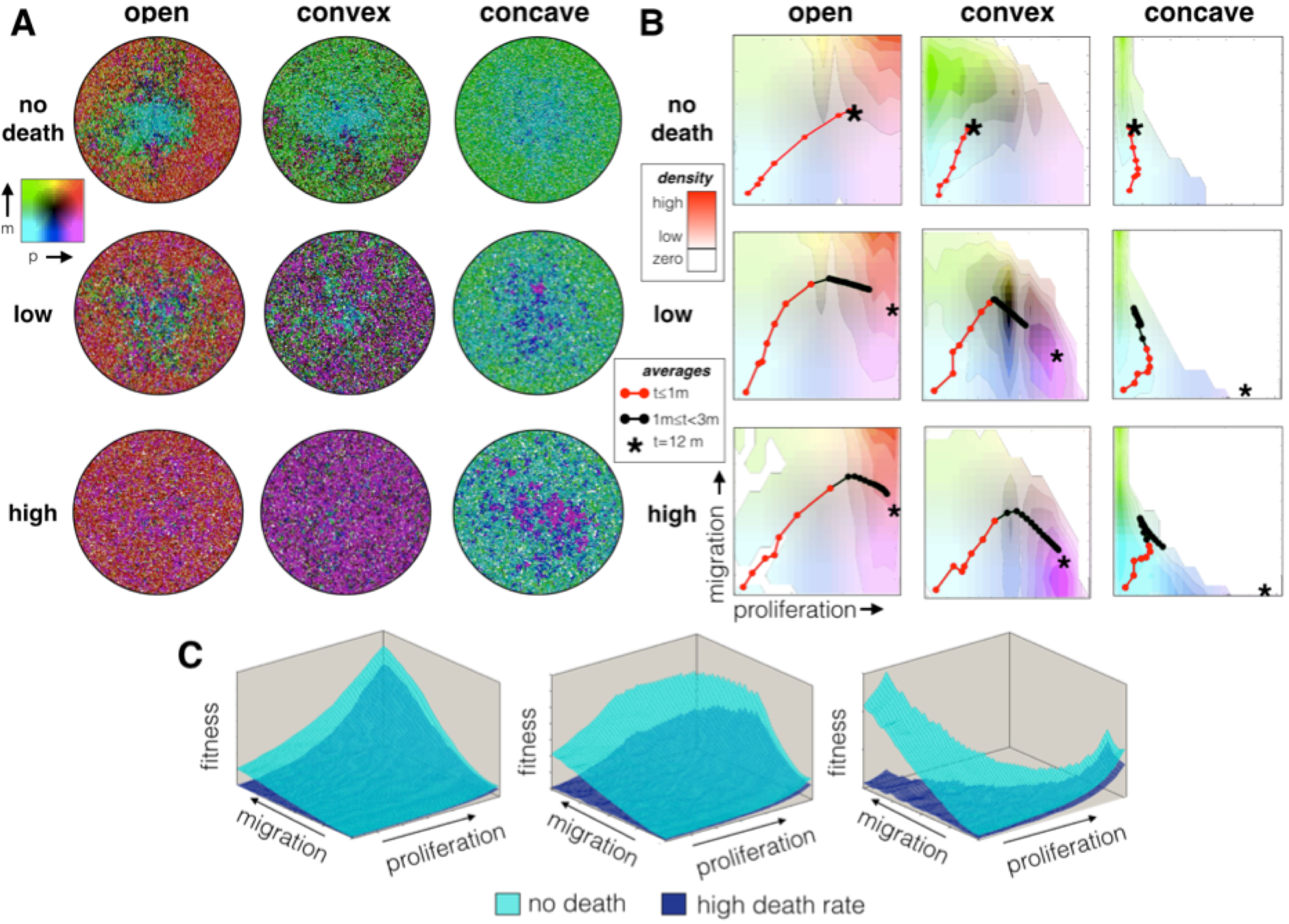
The effects of the death rate (no death, low, and high) and tradeoff boundaries (open, convex and concave) on the evolution of migration and proliferation rates. The probability of death for a single cell is once per week (high death rate) and once every two weeks (low death rate). A) The spatial layout is shown and B) and the frequency of trait combinations is shown, where the red points and line mark the average trait values every 5 days for the first month. The black points show the continued evolutionary trajectory up until 3 months, and the asterisk, the final value at 12 months. The background colors correspond to the density of cell traits after reaching capacity; Brightly colored areas correspond to high densities, and the completely white area contains no cells. C) Increasing the death rate reduces the fitness (growth rate) of a tumor population. The tradeoff boundary affects which traits are most fit.

When the cells have completely filled the space, the shape of the phenotypic tradeoff boundary (open, convex or concave) strongly influences the endpoint of evolution. Regardless of the death rate, the open boundary favors fast proliferation and high migration speeds. However, as space fills migration speeds matter much less than proliferation rates. Mutational drift in the migration trait leads to a lower mean and higher phenotypic variance than seen in the proliferation trait. Phenotypes with moderate to low proliferation rates become absent with time. With a low death rate, the convex boundary favors the coexistence of a more generalist phenotype with a fast proliferation phenotype. With a low death rate, the convex boundary sees the generalist phenotype outcompeted by fast proliferating cells with lower migration rates. With a low death rate, the concave boundary, as predicted, favors the coexistence between cancer cells with fast proliferation (but low migration) and cells with fast migration (but low proliferation). With a high death rate, the concave boundary favors cancer cells with high proliferation rates at the expense of migration. Comparing the fitness landscapes for each trade-off for high death rate and no death shows peaks where each of these phenotypes are favored (Fig. 4C).

### SPATIALLY CLUSTERED DEATH CATASTROPHES SELECT FOR MIGRATION

We introduced significant environmental stochasticity by having all individuals die within a randomly selected 500 μm diameter circular area. This regional catastrophe might represent a sudden (and temporary) loss of blood vasculature, immune cell intrusion, or pooling of toxic metabolites. While keeping the probability of death constant at one death per week per cell (high death rate) we compared three mortality regimes where we varied the fraction of deaths occurring by demographic stochasticity (random cell death) relative to environmental stochasticity (catastrophes). The three regimens had 0%, 50% and 100% catastrophic death. The results are shown in Fig. 5.

**Figure 5.**
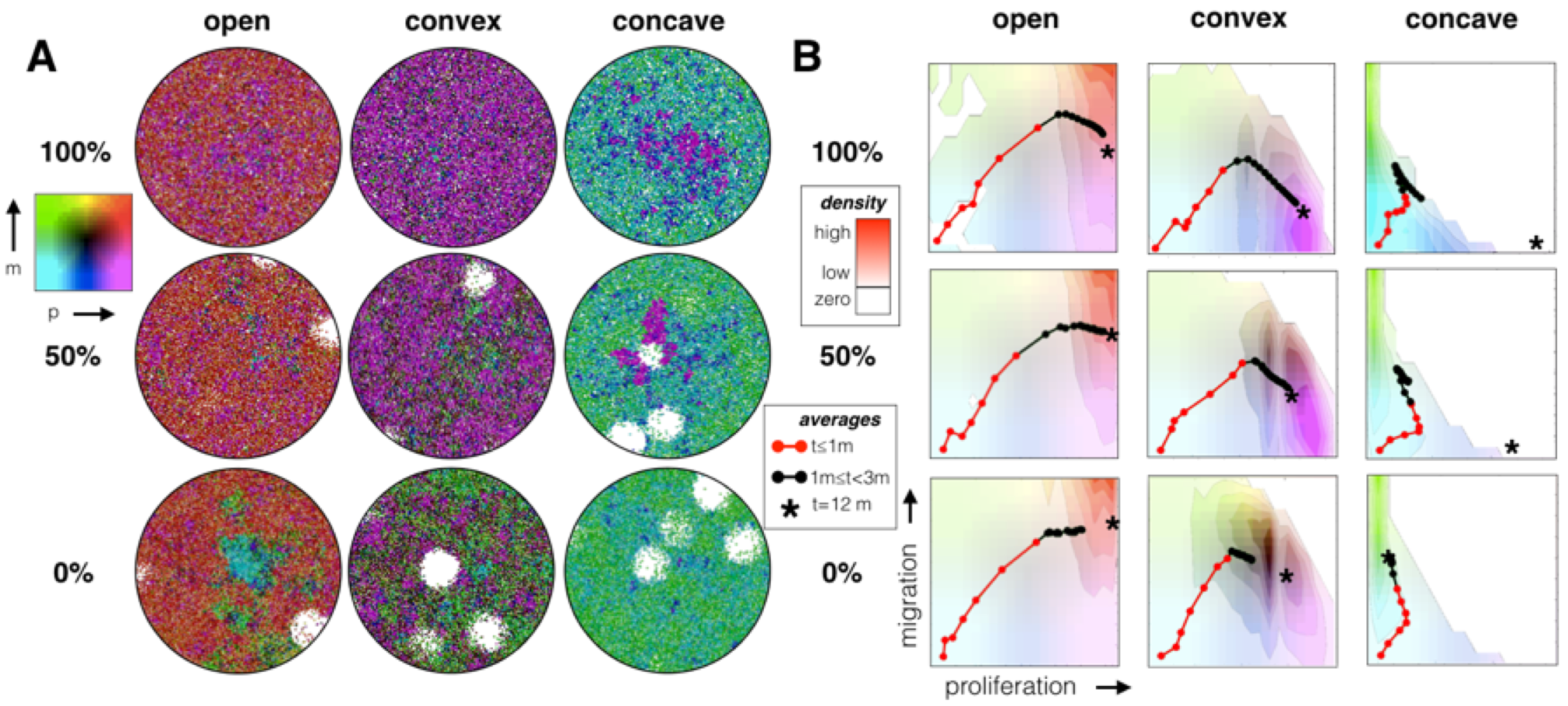
The percent of death that is random vs catastrophic is varied. The top row has 0% catastrophic and 100% random death, the middle row, 50% catastrophic and 50% random death, and the bottom row, 100% catastrophic and 0% random death. The death rate is once per week per cell (same as the high death rate from Fig. 4). A) The spatial layout is shown and B) and the frequency of trait combinations is shown, where the red points and line mark the average trait values every 5 days for the first month. The black points show the continued evolutionary trajectory up until 3 months, and the asterisk, the final value at 12 months. The background colors correspond to the density of cell traits after reaching capacity; Brightly colored areas correspond to high densities, and the completely white area contains no cells.

In all cases raising the percent of deaths by catastrophes increases selection for migration. For the open tradeoff boundary, this results in a similarly high proliferation rate even as the migration rate increases with environmental stochasticity. For the convex tradeoff, there is more variance in phenotypic properties. But, as environmental stochasticity increases, migration is favored over proliferation with a very generalist phenotype emerging when all deaths are catastrophic. For the concave tradeoff boundary, the average phenotype switches from high proliferation and low migration to low proliferation and high migration as environmental stochasticity goes from 0% to 100% of the cause of death.

## IV. DISCUSSION

Rates of cell turnover matter. As expected, in our model, increasing the death rate speeds the rate of evolution while having little impact on the endpoint of evolution or the equilibrium population size of cancer cells at the end of the simulation (12 months). Our off-lattice model places an upper bound on the space available for cells. While increasing the death rate opens up space, cells fill it quickly as neighboring cells now have the opportunity to successfully proliferate (at even the lowest proliferation rate cells divide once every 50 hours). Longer runs of cell proliferation permit the accumulation of mutations that can increase migration and/or proliferation rates. However, with no deaths, evolution eventually stops on the interior of the tumor and can only occur along the expanding boundary. One sees concentric rings of more highly adapted cells as we move from the center to the edge of the tumor. This is not the case when there is continual cell turnover. While slower in the interior than edge of the tumor, evolution proceeds with the replacement of less fit individuals by those with either higher combined rates of proliferation and migration, or individuals with more successful combinations of traits when the trait-tradeoff boundary has been reached.

The results illustrate the direct impact of cell turnover, throughout the habitable regions, on tumor evolutionary dynamics. However, not all ecological and evolutionary models in the literature incorporate cell death and cell replacement. The distribution of phenotypes among cancer cells in a tumor represent a balance between mutation, drift and selection. With each cell division, mutations can occur that randomly alter proliferation and migration. Those generating higher fitness should increase in frequency, but a large amount of heritable variation is maintained within the tumor due to the stochastic nature of births, deaths and mutations; the lower the rate of cell turnover, the higher the phenotypic variability among cancer cells. In reality, tumors exhibit large amounts of genetic variation - the extent to which this is maintained by mutation and drift and purged through selection remains an open and important question (17,30–32).

Furthermore the edge of the tumor likely offers very different conditions in terms of substrates, normal cell architecture and exposure to the immune system (33,34). Hence, a number of agent-based models focus on tumor spatial heterogeneity in an environmental context, such as normal cells, stroma, and vasculature (4,5,35,36). Here, we considered a much simpler model where all space is equal without regard for the position of blood vessels, and the only factor creating heterogeneity is the number and dispersion of cancer cells. This means our model has two rather distinct phases. During the first, natural selection favors migration over proliferation as the tumor expands into pristine space, the second favors proliferation over migration once the space has been filled by the cancer cells. This accords with the observation that the edge of tumors may select for more “aggressive” cancer cells defined as those more likely to migrate, invade surrounding tissue, and perhaps initiate metastases (37,38).

There are direct parallels, of our results to ecological systems in which mortality can take the form of the stochastic death of an individual (demographic stochasticity) or the catastrophic death of a group of individuals (environmental stochasticity). In forests, for instance, individual deaths of trees create small gaps in the canopy whereas the blowdown of a group of neighboring trees create large gaps. The size and nature of gaps can result in the slower or faster regeneration of different tree species (39). Our model considers the eco-evolutionary consequences of different size gaps in the tumor created by either demographic or environmental stochasticity (while holding overall mortality rates constant). As seen in many natural systems, small gaps select for proliferation over dispersal and vice-versa for large gaps (40). While understudied, temporal variability in local blood flow, immune intrusion, hypoxia, and Ph likely result in varying degrees of local and catastrophic mortality followed by opportunities for recolonization. Generally, histology from biopsies or radiographic imaging of tumors produce a static snapshot that cannot track the fates of individual cells within small regions of a tumor. Our model provides a platform to study how death affects the competition of cells for space and their subsequent evolution.

In the absence of tradeoffs, natural selection should favor improvements in all traits that enhance fitness. Several lines of evidence empirically suggest tradeoffs between the traits proliferation versus migration or “go-or-grow” models (9,23,24). As expected in our model, the lack of a tradeoff saw rapid increases in both proliferation and migration. Although with demographic stochasticity and filled space, migration is no longer under strong selection. At this point, mutation and drift create a lower mean migration rate with large phenotypic variance. Generally, a convex tradeoff selected for a generalist phenotype. Under demographic stochasticity this resulted in high phenotypic variance and sometimes the dominance of high proliferation, low migration phenotypes. Environmental stochasticity selected for the more generalist phenotype. A concave phenotypic tradeoff boundary always selected for low proliferation, high migration phenotypes during the tumor’s expansion phase. These were then replaced by high proliferation, low migration phenotypes once the tumor achieved maximum size. The only exception occurred with environmental stochasticity where the high migration phenotype continues to be favored. Just as with experiments that determine the cost of resistance in cancer cells (41–44), establishing the nature of tradeoffs between migration and proliferation may require cell culture experiments where the cancer cells are grown under conditions of strong nutrient limitation. Tradeoffs between dispersal and survival or fecundity and dispersal are common in natural plant and animal species (45,46).

In our model, we assume death rates are independent of phenotype. Thus, natural selection favors phenotypes that maximize the probability of cell division. This probability represents fitness *f*, and it is the product of the proliferation rate *p* and the probability of having space around the cell to proliferate s: *f*=*sp*. We can see the direct effect of increasing the proliferation rate on fitness by taking the derivative of fitness with respect to the proliferation rate (*df*/*dp*=*s*). Likewise, the effect on fitness of increasing the migration rate, *m*, can be found by taking its derivative with respect to the migration speed *df/dm*. This depends not only on the proliferation rate but also how much space is opened up by moving: *df/dm*=*p*\**ds*/*dm*. Some of our observations can be understood through these relationships. During the expansion phase of the tumor, *s* is relatively large and the space gained by increasing migration, *ds/dm*, is very large. Selection will be strong for both proliferation and migration but relatively stronger for migration. When the space in the tumor core is completely filled both *s* and *ds/dm* go to zero. Hence there is no longer positive selection for migration. By creating many small gaps, demographic stochasticity creates space and thus maintains selection for proliferation. Because the spaces are small, there are no benefits to migration. Environmental stochasticity creates the same amount of total space over time as demographic stochasticity but this space is more contiguous. Migration is favored as a means of exploiting empty regions. Thus, *ds/dm* will be larger and positive. There will be positive selection for both proliferative and migratory phenotypes, but a larger selection for migratory phenotypes. Some of these properties will be general to all organisms (e.g., cane toads (47), and house sparrows spreading in Kenya (48)), and others just to cancer because it is a densely packed, asexual, and single-celled organism.

Our model has similarities to other models and systems where selection balances two traits. In natural systems this can take the form of seed dispersal versus dormancy in annual plants where the former transports the individual spatially and the latter temporally to less crowded and more favorable places (49,50). In dispersal-dormancy models, the traits may exhibit tradeoffs via seed size, seed coat thickness, and features that enhance dispersal such as burrs and samara (wings). In cancer, a number of agent-based models consider vector-valued traits. These include degree of glycolytic respiration (Warburg effect) and tolerance to acidic conditions. While not necessarily linked through tradeoffs, the two traits become co-adapted as increased glycolysis promotes acidic conditions necessitating the evolution of acid tolerance (5). Spatial models often see rings of different trait values extending from the interior to the tumor’s boundary (3,4,6). In these models, selection happens solely at the tumor edge where there is space to proliferate. In relation to these works our model invites spatially-explicit investigations into how traits evolve in response to population spread, death rates, and demographic versus environmental stochasticity. It emphasizes the critical need to estimate cell turnover via measurements of both death and proliferation rates. A variety of markers and metrics exist for measuring proliferation (e.g. Ki67, mitotic index) and death (e.g. caspases, TUNEL assay). However, these are often just surrogates, rarely measured simultaneously, and generally cannot be measured in vivo. Because of these challenges, most data simply describe net tumor growth (i.e. doubling times). We advocate deconstructing this net metric into distinct fractions of proliferating, quiescent, and dying. To study evolving traits such as proliferation-migration trade-offs, we see a need for non-destructive sensors/markers of cell processes that can be measured through both space and time.

## ACKNOWLEDGMENTS

The authors gratefully acknowledge Mehdi Damaghi for the tumor spheroid model graphic (Fig. 1A) and Mark Lloyd for the digital pathology graphic (Fig. 1B). Funding came from both the Cancer Systems Biology Consortium (CSBC) and the Physical Sciences Oncology Network (PSON) at the National Cancer Institute, through grants U01CA151924 (supporting A. Anderson and J. Gallaher) and U54CA193489 (supporting A. Anderson and J. Brown).

## REFERENCES

1. Wallace D, Guo X. Properties of tumor spheroid growth exhibited by simple mathematical models. Frontiers in oncology. 2013.

2. Anderson ARA, Weaver AM, Cummings PT, Quaranta V. Tumor morphology and phenotypic evolution driven by selective pressure from the microenvironment. Cell. 2006 Dec 1;127(5):905–15.

3. Gallaher JA, Anderson ARA. Evolution of intratumoral phenotypic heterogeneity: the role of trait inheritance. 2013 Aug 6;3(4):20130016.

4. Frankenstein Z, Basanta D, Franco OE, Gao Y, Javier RA, Strand DW, et al. Stromal Reactivity Differentially Drives Tumor Cell Evolution and Prostate Cancer Progression. 2017 Jul 4;:1–46.

5. Robertson-Tessi M, Gillies RJ, Gatenby RA, Anderson ARA. Impact of metabolic heterogeneity on tumor growth, invasion, and treatment outcomes. Cancer Research. 2015 Apr 15;75(8): 1567–79.

6. Poleszczuk J, Hahnfeldt P, Enderling H. Evolution and Phenotypic Selection of Cancer Stem Cells. Rzhetsky A, editor. PLoS Computational Biology. 2015 Mar 5;11(3):e1004025–14.

7. Waclaw B, Bozic I, Pittman ME, Hruban RH, Vogelstein B, Nowak MA. A spatial model predicts that dispersal and cell turnover limit intratumour heterogeneity. Nature. 2015 Aug 26;525(7568):261–4.

8. Bozic I, Allen B, Nowak MA. Dynamics of targeted cancer therapy. Trends in Molecular Medicine. 2012 Jun;18(6):311–6.

9. Kaznatcheev A, Scott JG, Basanta D. Edge effects in game-theoretic dynamics of spatially structured tumours. Journal of The Royal Society Interface. 2015 Jun 3;12(108):20150154–4.

10. Kerr JFR, Wyllie AH, Currie AR. Apoptosis: A Basic Biological Phenomenon with Wide-ranging Implications in Tissue Kinetics. British Journal of Cancer. Nature Publishing Group; 1972 Aug 1;26(4):239–57.

11. Alenzi FQB. Links between apoptosis, proliferation and the cell cycle. British Journal of Biomedical Science. 2016 May 23;61(2):99–102.

12. Liu S, Edgerton SM, Moore DH, Thor AD. Measures of cell turnover (proliferation and apoptosis) and their association with survival in breast cancer. Clinical Cancer Research. 2001 Jun;7(6):1716–23.

13. Soini Y, Pääkkö P, Lehto VP. Histopathological evaluation of apoptosis in cancer. The American Journal of Pathology. 1998 Oct;153(4):1041–53.

14. Zimmerman MA, Huang Q, Li F, Liu X, Li C-Y. Cell death-stimulated cell proliferation: a tissue regeneration mechanism usurped by tumors during radiotherapy. Seminars in Radiation Oncology. 2013 Oct;23(4):288–95.

15. Labi V, Erlacher M. How cell death shapes cancer. Cell Death Dis. 2015 Mar 5;6(3):e1675.

16. Hamilton WD, May RM. Dispersal in stable habitats. Nature. 1977.

17. Sottoriva A, Kang H, Ma Z, Graham TA, Salomon MP. A Big Bang model of human colorectal tumor growth. Nature. 2015.

18. Bozic I, Gerold JM, Nowak MA. Quantifying Clonal and Subclonal Passenger Mutations in Cancer Evolution. PLoS Computational Biology. 2016 Feb;12(2):e1004731.

19. Aktipis CA, Boddy AM, Gatenby RA, Brown JS, Maley CC. Life history trade-offs in cancer evolution. Nat Rev Cancer. Nature Publishing Group; 2013 Nov 11;13(12):883–92.

20. Shoval O, Sheftel H, Shinar G, Hart Y, Ramote O, Mayo A, et al. Evolutionary Trade-Offs, Pareto Optimality, and the Geometry of Phenotype Space. Science. 2012 May 31;336(6085):1157–60.

21. Orlando PA, Gatenby RA, Brown JS. Tumor evolution in space: the effects of competition colonization tradeoffs on tumor invasion dynamics. Frontiers in Oncology. 2013;3:45.

22. Gerlee P, Nelander S. The impact of phenotypic switching on glioblastoma growth and invasion. Alber MS, editor. PLoS Computational Biology. 2012;8(6):e1002556.

23. Hatzikirou H, Basanta D, Simon M, Schaller K, Deutsch A. “Go or grow”: the key to the emergence of invasion in tumour progression? Math Med Biol. 2012 Mar;29(1):49–65.

24. Gerlee P, Anderson ARA. Evolution of cell motility in an individual-based model of tumour growth. Journal of Theoretical Biology. 2009 Jul;259(1):67–83.

25. Levins R. Evolution in changing environments: some theoretical explorations. 1968.

26. Gatenby RA, Cunningham JJ, Brown JS. Evolutionary triage governs fitness in driver and passenger mutations and suggests targeting never mutations. Nature Communications. 2014.

27. Engen S, Bakke Ø, Islam A. Demographic and environmental stochasticity-concepts and definitions. Biometrics. 1998 Sep 1;54(3):840–6.

28. Gelles JD, Chipuk JE. Robust high-throughput kinetic analysis of apoptosis with real-time high-content live-cell imaging. Cell Death Dis. 2016.

29. van Nieuwenhuijze AEM, Van Lopik T, Smeenk RJT, Aarden LA. Time between onset of apoptosis and release of nucleosomes from apoptotic cells: putative implications for systemic lupus erythematosus. Annals of the Rheumatic Diseases. BMJ Publishing Group Ltd; 2003 Jan 1;62(1): 10–4.

30. Iwasaki WM, Innan H. Simulation framework for generating intratumor heterogeneity patterns in a cancer cell population. PLoS ONE. 2017;12(9):e0184229.

31. Durrett R, Foo J, Leder K, Mayberry J, Michor F. Intratumor heterogeneity in evolutionary models of tumor progression. Genetics. 2011 Jun;188(2):461–77.

32. Gerlinger M, Rowan A, Horswell S. Intratumor Heterogeneity and Branched Evolution Revealed by Multiregion Sequencing. New England Journal of Medicine. 2012.

33. Ramamonjisoa N, Ackerstaff E. Characterization of the Tumor Microenvironment and Tumor-Stroma Interaction by Non-invasive Preclinical Imaging. Frontiers in Oncology. 2017;7:3.

34. Lloyd MC, Cunningham JJ, Bui MM, Gillies RJ, Brown JS, Gatenby RA. Darwinian Dynamics of Intratumoral Heterogeneity: Not Solely Random Mutations but Also Variable Environmental Selection Forces. Cancer Research. 2016 Jun 1;76(11):3136–44.

35. Ibrahim Hashim A, Robertson-Tessi M, Enriquez-Navas PM, Damaghi M, Balagurunathan Y, Wojtkowiak JW, et al. Defining Cancer Subpopulations by Adaptive Strategies Rather Than Molecular Properties Provides Novel Insights into Intratumoral Evolution. Cancer Research. American Association for Cancer Research; 2017 May 1;77(9):2242–54.

36. Basanta D, Anderson ARA. Homeostasis Back and Forth: An Eco-Evolutionary Perspective of Cancer. 2016 Dec 6;:1–23.

37. Clark AG, Vignjevic DM. Modes of cancer cell invasion and the role of the microenvironment. Current Opinion in Cell Biology. 2015 Oct;36:13–22.

38. Petrie RJ, Yamada KM. At the leading edge of three-dimensional cell migration. Journal of Cell Science. 2013 Feb 27;125(24):5917–26.

39. Canham CD. Different responses to gaps among shade-tolerant tree species. Ecology. 1989;70(3):548–50.

40. Nagel TA, Svoboda M, Kobal M. Disturbance, life history traits, and dynamics in an old-growth forest landscape of southeastern Europe. Ecol Appl. 2014 Jun;24(4):663–79.

41. Chmielecki J, Foo J, Oxnard GR, Hutchinson K, Ohashi K, Somwar R, et al. Optimization of dosing for EGFR-mutant non-small cell lung cancer with evolutionary cancer modeling. Science Translational Medicine. American Association for the Advancement of Science; 2011 Jul 6;3(90):90ra59–9.

42. Moore N, Houghton JM, Lyle S. Slow-cycling therapy-resistant cancer cells. Stem cells and development. 2011.

43. Silva AS, Kam Y, Khin ZP, Minton SE, Gillies RJ, Gatenby RA. Evolutionary Approaches to Prolong Progression-Free Survival in Breast Cancer. Cancer Research. 2012 Dec 16;72(24):6362–70.

44. Kreso A, O’Brien CA, van Galen P, Gan OI, Notta F, Brown AMK, et al. Variable Clonal Repopulation Dynamics Influence Chemotherapy Response in Colorectal Cancer. Science. 2013 Jan 31;339(6119):543–8.

45. Duthie AB, Abbott KC, Nason JD. Trade-offs and coexistence in fluctuating environments: evidence for a key dispersal-fecundity trade-off in five nonpollinating fig wasps. The American Naturalist. 2015 Jul;186(1):151–8.

46. Weigang HC, Kisdi É. Evolution of dispersal under a fecundity-dispersal trade-off. Journal of Theoretical Biology. 2015 Apr;371:145–53.

47. Phillips BL, Brown GP, Greenlees M, Webb JK, Shine R. Rapid expansion of the cane toad (Bufo marinus) invasion front in tropical Australia. Austral Ecol. Blackwell Publishing Asia; 2007 Apr;32(2):169–76.

48. Schrey AW, Liebl AL, Richards CL, Martin LB. Range expansion of house sparrows (Passer domesticus) in Kenya: evidence of genetic admixture and human-mediated dispersal. J Hered. 2014 Jan;105(1):60–9.

49. Venable DL, Brown JS. The selective interactions of dispersal, dormancy, and seed size as adaptations for reducing risk in variable environments. The American Naturalist. 1988;131(3):360–84.

50. Gremer JR, Venable DL. Bet hedging in desert winter annual plants: optimal germination strategies in a variable environment. Ecol Lett. 2014 Mar;17(3):380–7.

